# Hierarchy of spindle forces in prometaphase

**DOI:** 10.64898/2026.10.28.762694

**Authors:** Wenzheng Shi, Fioranna Renda, Alexey Khodjakov, Christopher E. Miles, Alex Mogilner

## Abstract

Dramatic movements characterize rapid assembly of mitotic spindle in prometaphase: chromosomes congress to the spindle equator, while spindle poles segregate rapidly elongating the spindle. Direct biophysical measurements of forces driving these movements are lacking, but the forces could be learned from the rapid pole and chromosome movements. We analyzed 3D trajectories of chromosomes and spindle poles in the wild-type and PRC1-depleted cells during early prometaphase and reconstructed the spindle forces. This analysis posits that (i) chromosomes do not interact with each other, except for a short-range steric repulsion, (ii) effective pole-chromosome forces are repulsive at short and attractive at long distances, (iii) dominant forces of spindle poles segregation are interpolar in origin. The important implication of these results is that the pole-pole segregation is robust, largely insensitive to regional variations inside the cell and to complex process of chromosome congression.

**SIGNIFICANCE:** Mechanics of prometaphase is crucial for mitotic spindle assembly, yet measurements of forces in prometaphase are lacking. We use movements of centrosomes and chromosomes in prometaphase to reconstruct the spindle forces. We find that the forces are hierarchical, with the interpolar force balance dominating smaller forces between the spindle poles and chromosomes. The important implication of these results is that the pole-pole segregation is robust, largely insensitive to regional variations inside the cell and to complex process of chromosome congression.

## INTRODUCTION

Mitotic spindle, a molecular machine segregating chromosomes at the onset of cell division in eukaryotic cells, self-assembles from microtubules (MTs), chromosomes (CHs) and molecular motors in early stages of mitosis (1). One of the most important, and yet incompletely understood, of these stages is prometaphase (2). In animal somatic cells, the spindle assembles when MTs connect CHs to two centrosomes (future spindle poles) after nuclear envelope breakdown. In early prometaphase, as these connections are made, interpolar MT bundles assemble making the spindle (3, 4), while CHs congress to the spindle surface (5, 6) and the spindle poles rapidly segregate (Fig. 1A, B, and D).

**Figure 1:**
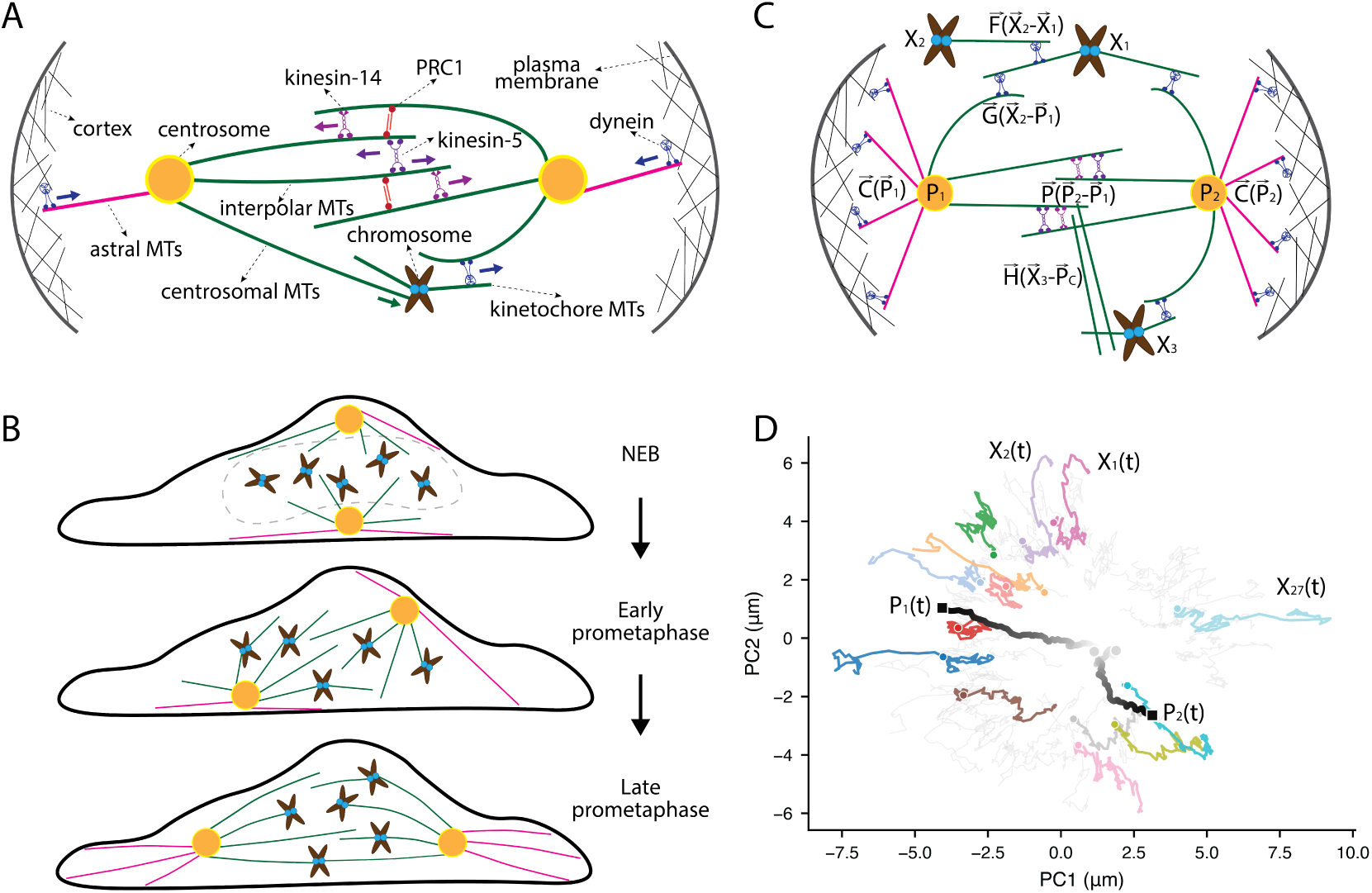
Spindle forces and movements in early prometaphase. **(A)** Schematic of the principal force-generating elements in prometaphase. Astral microtubules (MTs) link each centrosome to the cell cortex, where dynein pulls on them. Centrosomal MTs push on chromosome (CH) arms directly and through chromokinesin action. Dynein motors on short kinetochore-associated MTs pull CHs toward the centrosomes on long centrosomal MTs. Antiparallel interpolar MTs from the two centrosomes overlap between the poles; on the overlaps, stabilized by PRC1, bipolar kinesin-5 motors slide the MTs apart segregating the spindle poles, while kinesin-14 motors pull the MTs together resisting the kinesin-5 action. **(B)** Schematic side view of an RPE-1 cell at nuclear envelope breakdown (NEB, top), early prometaphase (middle) and late prometaphase (bottom). In a majority of cells, at NEB, the centrosomes are at the top and bottom of the flattened nucleus (dashed). During early prometaphase, the centrosomes segregate and move toward opposite sides of the former nucleus, rotating the pole–pole axis toward the substrate plane. By late prometaphase the poles lie on opposite sides of the cell and the CHs have congressed between them. **(C)** Effective particle model (Eqs. 1–2). CHs (*X_i_*) and centrosomes (*P*_1_, *P*_2_) interact through pairwise distance-dependent forces: *F*, between pairs of CHs; *G*, between a CH and a centrosome; *H*, between a CH and the spindle center *P_c_* (the midpoint between the two centrosomes); *P*, between the two centrosomes (net interpolar force); and *C*, between each centrosome and the cortex (net astral force). **(D)** Measured trajectories in a representative control cell, projected onto the first two principal components (PC1, PC2) of that cell’s pooled centrosome and CH positions, i.e. a single time-independent, per-cell basis computed from all poles, all CHs and all frames together. PC1 runs close to the pole–pole axis, so the projection shows pole separation and CH congression in one plane (see Methods). Centrosome trajectories (*P*_1_ (*t*), *P*_2_ (*t*)) are shaded from light to dark grey with time; circles mark the start and squares the end. A subset of chromosome trajectories (*X_i_* (*t*)) is shown with one color per chromosome (color does not encode time); the remaining chromosomes are shown in light grey.

Forces and movements within the spindle are central to understanding how the high fidelity of chromosome segregation is achieved (7, 8). These forces and movements are primarily generated by a host of MT-based molecular motors (Fig. 1A). First, dyneins pulling on astral MTs (9–11) contribute to movements of two centrosomes (CSs) toward opposite parts of cell cortex causing segregation of CSs and increase in spindle length (12). Dyneins also pull short MTs attached to kinetochores toward minus ends of the long centrosomal MTs (Fig. 1A), effectively pulling CHs toward the CSs (5, 6). Other centrosomal MTs contact CH arms, pushing CHs away from the CSs through both polymerization force and actions of specialized motors (13–15). Last, but not least, a set of antiparallel interpolar MTs overlap in the middle between the spindle poles and get connected, among other motors and crosslinkers, by kinesin-5 and kinesin-14 (16) (Fig. 1A). Bipolar kinesin-5 motor slides the interpolar MTs apart effectively pushing the poles away from each other, while kinesin-14 motors bind to one of the MTs and pulls it toward the pole-located minus end of another MT effectively pulling the poles together. The resulting tug-of-war between these motors generates a net interpolar force. For this tug-of-war to happen, crosslinkers like PRC1 must be present in sufficient numbers to stabilize antiparallel interpolar MT bundles (17). Other processes, including, but not restricted to, cortex flows effectively pulling on astral MTs (18) and complex dynamic connections between different sets of MTs (19) contribute to these motor-generated forces.

Very few direct force measurements in the spindle were made. Landmark 1983 study by R.B. Nicklas measured a force to stop a CH during anaphase in grasshopper spermatocytes by using a calibrated micro-needle and found a surprisingly high force of hundreds of pN per CH (20). Magnetic tweezers were used to measure a ∼5 pN force of dynein-powered astral MT pull (11). A whole-spindle mechanics was probed revealing viscoelastic properties of the metaphase spindle (21). These measurements suggest that on the time scale of tens of seconds, the spindle behaves like a viscous liquid (21, 22).

This lack of direct mechanical measurements was partially compensated for by computational modeling of changes in spindle dynamics observed under various experimental perturbations. Around 20 astral MTs are predicted to pull on each CS (23, 24), which, considering the 5 pN-range force per astral MT (11) means ∼100 pN astral outward force per pole (24). Effective spindle pole viscous drag coefficient of hundreds of pN s *μ*m^−1^ is estimated (25). Considering characteristic cytoplasmic viscosity and observed rates of movements, an average net force on a CH should be on the pN scale (25). Similar pN-scale effective repulsion force between the CS and a CH arm was posited in (25, 26). Considering tens of CHs in the spindle (for example, 46 CHs in human cells), net force of tens of pN is proposed to be associated with CS-CH interactions (24). Rough estimates in (27) suggest that hundreds to thousands of kinesin-5 and kinesin-14 motors per spindle interact with the interpolar MT overlaps, which amounts to hundreds to thousands pN of the opposing forces from these two motors balancing each other on the interpolar MTs (16). These estimates were made based on data from a gamut of cell types and experimental assays, which make these numbers no more than a useful starting point.

In late prometaphase, the spindle almost reaches a mechanical steady state in making spindle responses to perturbations easier to interpret. This notion was used in (16, 24) to analyze motor forces and numbers in late prometaphase spindle in Drosophila melanogaster embryos. However, large movements of CSs and CHs in earlier prometaphase offer an advantage of a nonperturbative, observational, approach: inferring spindle forces from measured movements. This approach – reverse-engineering of interaction forces from measured trajectories (28–31) and from noise analysis of these trajectories (23) – has long history of success in cell biophysics.

In this paper, we use experimental data (reported in (5, 6)) from high-resolution 3D tracking of all CSs and CHs in human RPE1 cells (Fig. 1D) to infer spindle forces (Fig. 1C) in early prometaphase. In majority of these cells, two CSs start at the dorsal and ventral surfaces of the flattened nucleus (Fig. 1B), the envelope of which is breaking apart at the prometaphase onset. (Molecular processes during prophase leading to this prometaphase initial condition remain obscure (32, 33). Therefore, they are not within the scope of the analyses presented here.) Then, the CSs move apart to the sides of the former nucleus, while CHs congress toward the spindle. As quantitative information is lacking on when and where individual MTs and motors interact, it is useful to consider the CS-CH system as a ‘particle’ system (28): 2 CSs and 46 CHs interact with effective pairwise distance-dependent forces (see Methods). What are the relative magnitudes of interpolar vs astral vs CS-CH forces? What are distance dependencies of these forces? Do CHs interact mechanically with each other?

Our analyses of the particle models suggest that CHs only interact with each other through a short-range steric repulsion, that effective CS-CH forces are repulsive at short (up to ∼4 *μ*m) and attractive at longer distances, and that dominant repulsive force of spindle poles segregation is interpolar in origin and is generated by hundreds of counterbalanced kinesin-5 and kinesin-14 motors, assisted little by astral MT forces and by net repulsive CS-CH forces. The resulting model explains rapid, steady, and robust rate of spindle elongation during early prometaphase.

## MATERIALS AND METHODS

### Trajectory data

We analyzed three-dimensional trajectories of the two centrosomes and the chromosomes in human RPE-1 cells, followed in three-dimensional space at 5 s intervals and reported previously (5, 6). The dataset comprises 9 control cells and 12 cells depleted for PRC1 via siRNA. Interaction kernels were inferred from control cells, using the frames from NEB to roughly the end of pole separation (40% of the interval between NEB and anaphase onset). For display, the trajectories of each cell are projected onto the first two principal components of that cell’s pooled centrosome and chromosome positions (Figs. 1D and Fig. 3A, B).

### Effective particle model

Following (28, 29, 31), we treat the two centrosomes (spindle poles) and the chromosomes as particles that move overdamped under central, distance-dependent pairwise forces. Each force is written in velocity units, i.e., divided by the drag coefficient of the particle on which it acts. Positive kernel values denote repulsion.

#### Chromosomes

The position **x***_i_* of chromosome *i* evolves as

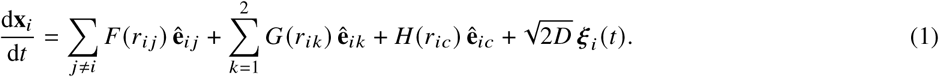

Here *r_ij_* = ||**x***_i_* – **x***_j_*||, *r_ik_* = ||**x***_i_* – **p***_k_*|| and *r_ic_* = ||**x***_i_* – **p***_c_*|| are the distances from chromosome *i* to chromosome *j*, to pole *k*, and to the spindle center **p***_c_* = (**p**_1_ + **p**_2_)/2, respectively. The **ê** are the corresponding unit vectors pointing from the partner toward chromosome *i*. *F* describes chromosome–chromosome interactions, for example steric exclusion or links through kinetochore-associated MTs or chromatin. *G* describes interactions with each pole through centrosomal MTs, and *H* an interaction with the spindle as a whole. The ***ξ****_i_* are independent three-dimensional Gaussian white noises, 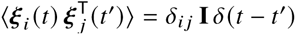, and *D* is an effective diffusivity. The pole trajectories **p***_k_*(*t*)are taken from the data and are not modeled in Eq. (1). All chromosomes obey the same law; differences between chromosomes, e.g. in size (34), are neglected.

We compared four topologies: (i) poles, with *G* ≠ 0 and *F* = *H* = 0; (ii) center, with *H* ≠ 0 and *F* = *G* = 0; (iii) poles + chromosomes (free), with *F*, *G* ≠ 0, *H* = 0 and *F* unconstrained; and (iv) poles + chromosomes (short range), as (iii) but with *F*(*r*) = *s*(*r*)*F̃* (*r*). The envelope 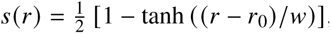, with *r*_0_ = 1.5 *μ*m and *w* = 0.3 *μ*m, restricts *F* to the contact range of chromosome-sized objects.

#### Poles

Pole *k* moves according to

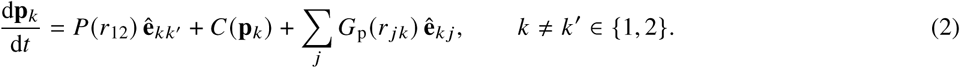

Here *r*_12_ is the pole–pole distance, **ê***_kk_*′ and **ê***_k j_* are unit vectors pointing toward pole *k* from the other pole and from chromosome *j*, and *r _jk_* = ||**x** *_j_* − **p***_k_*||. *P* is the net interpolar force and *C* the net astral force. *G*_p_ is the reaction of the chromosome–pole forces on the pole. By Newton’s third law, *G*_p_ = *cG*, where *G* is the kernel in Eq. (1) and *c* ≥ 0 is the ratio of the chromosome drag coefficient to the pole drag coefficient. The astral term *C* is omitted in all fits (see Results).

We compared three pole models: (i) interpolar only (pp), with *G*_p_ = 0; (ii) chromosome–pole only (cp-only), with *P* = 0 and *G*_p_ a free kernel; and (iii) a mixed model, with *P* free and *G*_p_ = *cG*, in which the shape of *G* is fixed to that of the short-range chromosome model and only the amplitude *c* is fitted.

In the mixed model, the interpolar and chromosome–pole contributions to the separation rate are nearly collinear in time, because both depend mainly on *r*_12_. The data therefore do not determine how the total force splits between them. We instead imposed the chromosome–pole fraction

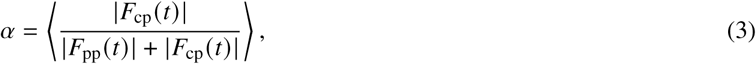

where *F*_pp_(*t*) and *F*_cp_(*t*) are the interpolar and chromosome–pole contributions to the pole-separation rate, and the average runs over frames and cells. For each target value of *α*, we minimized the loss in Eq. (4) subject to Eq. (3) and *c* ≥ 0, using sequential least-squares programming. The sign constraint excludes unphysical solutions in which *c* < 0 inverts the learned kernel.

### Kernel inference

Each kernel *f* (that is, *F̃*, *F*, *G*, *H*, *P* or *G*_p_) is expanded in *B* = 10 clamped cubic B-splines with uniformly spaced interior knots: 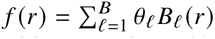. The drift, i.e. the deterministic right-hand side of Eq. (1) or Eq. (2), is then linear in the coefficients ***θ***. Velocities are estimated by forward differences, 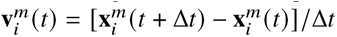, for particle *i* in cell *m*, with Δ*t* = 5 *s*. We minimize

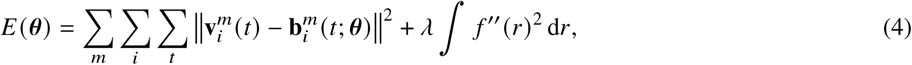

where 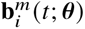 is the model drift evaluated at the observed positions at time *t*. The roughness penalty, with *λ* = 1, controls kernel smoothness. Minimizing Eq. (4) is a penalized linear least-squares problem. Its normal equations are assembled cell by cell and summed before a single solve (29, 30), which also makes leave-one-cell-out refits inexpensive.

#### Diffusivity and drift fraction

After the drift is fitted, the effective diffusivity is estimated from the residual displacements ***δ****_i_* (*t*) = **x***_i_* (*t* + Δ*t*) − **x***_i_* (*t*) − **b***_i_* (*t*) Δ*t* as *D* = ⟨∥***δ****_i_* ∥^2^⟩/(6Δ*t*). It is computed either pooled over all data (constant *D*) or as a smooth function *D d* of the distance *d* between the chromosome and the spindle center (Fig. 3C). To compare drift with diffusion over the elongation window *T* = 150 s, we compute

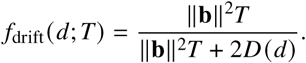

This is the fraction of the mean squared displacement along the drift direction, ||**b**||^2^*T*^2^ + 2*D*(*d*)*T*, that is due to drift. It is evaluated for each chromosome and frame and binned by *d* (Fig. 3D).

### Model validation and uncertainty

Models were compared by leave-one-cell-out cross-validation: each model was fit to all cells but one and evaluated on the held-out cell. For the CH topologies (Fig. 2C) and for the pp and cp-only pole models (Fig. 4D), the held-out error is the path error of a deterministic rollout: the noise-free model is integrated forward from the real initial frame using the fitted kernels, and the squared position error is averaged over all frames of the held-out cell. For the mixed pole model (Fig. 4F), in which the interpolar and CS–CH terms act together with the CS–CH kernel shape fixed to that learned from CH motion in all cells and their relative contribution set by the imposed share *α* (Eq. (3)), the held-out error is the root-mean-square error of the predicted pole-separation rate. Confidence bands on kernels are 5–95% percentile intervals over bootstrap.

**Figure 2:**
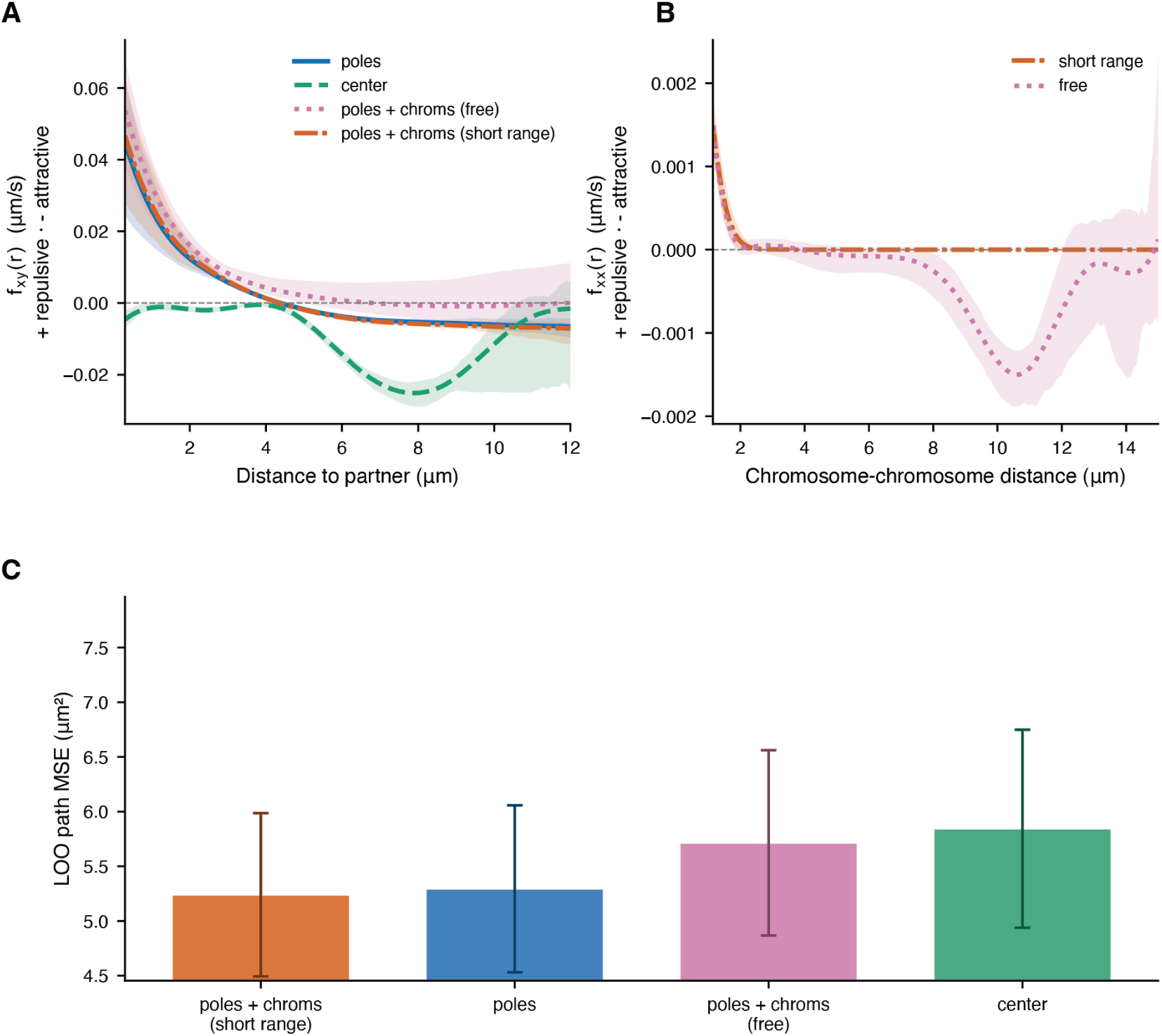
Chromosome–pole forces are repulsive at short range and attractive at long range; long-range chromosome–chromosome forces are not supported by the data. Four interaction topologies (Eq. 1) were fit to chromosome (CH) trajectories: CHs interacting with the poles only (poles; *G* ≠ 0); with the spindle midpoint only (*H* ≠ 0); with the poles and with each other through an unconstrained chromosome–chromosome kernel (poles + chroms, free; *G*, *F* ≠ 0); and with the poles and with each other through a short-range CH-CH kernel (poles + chroms, short range). Kernels are in velocity units (force divided by the effective drag coefficient); positive values denote repulsion and negative values attraction. **(A)** Chromosome–partner kernel *f_xy_* (*r*) as function of the distance *r* to the interaction partner (a pole for the three pole-based models; the spindle midpoint for the center model). **(B)** CH-CH kernel *f_xx_* (*r*) for the two topologies that include it. In the short-range model the kernel is multiplied by the steric envelope 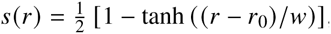, with *r*_0_ = 1.5 *μ*m and *w* = 0.3 *μ*m, which restricts it to the contact regime. In A and B, bands are 5–95% bootstrap confidence intervals. **(C)** Leave-one-cell-out path error: mean squared deviation between the observed chromosome positions in the held-out cell and a deterministic rollout of the fitted model (Eq. 1 without the noise term, driven by the measured pole trajectories). Bars show means over held-out cells. Models are ordered by mean value. Error bars show standard error.

**Figure 3:**
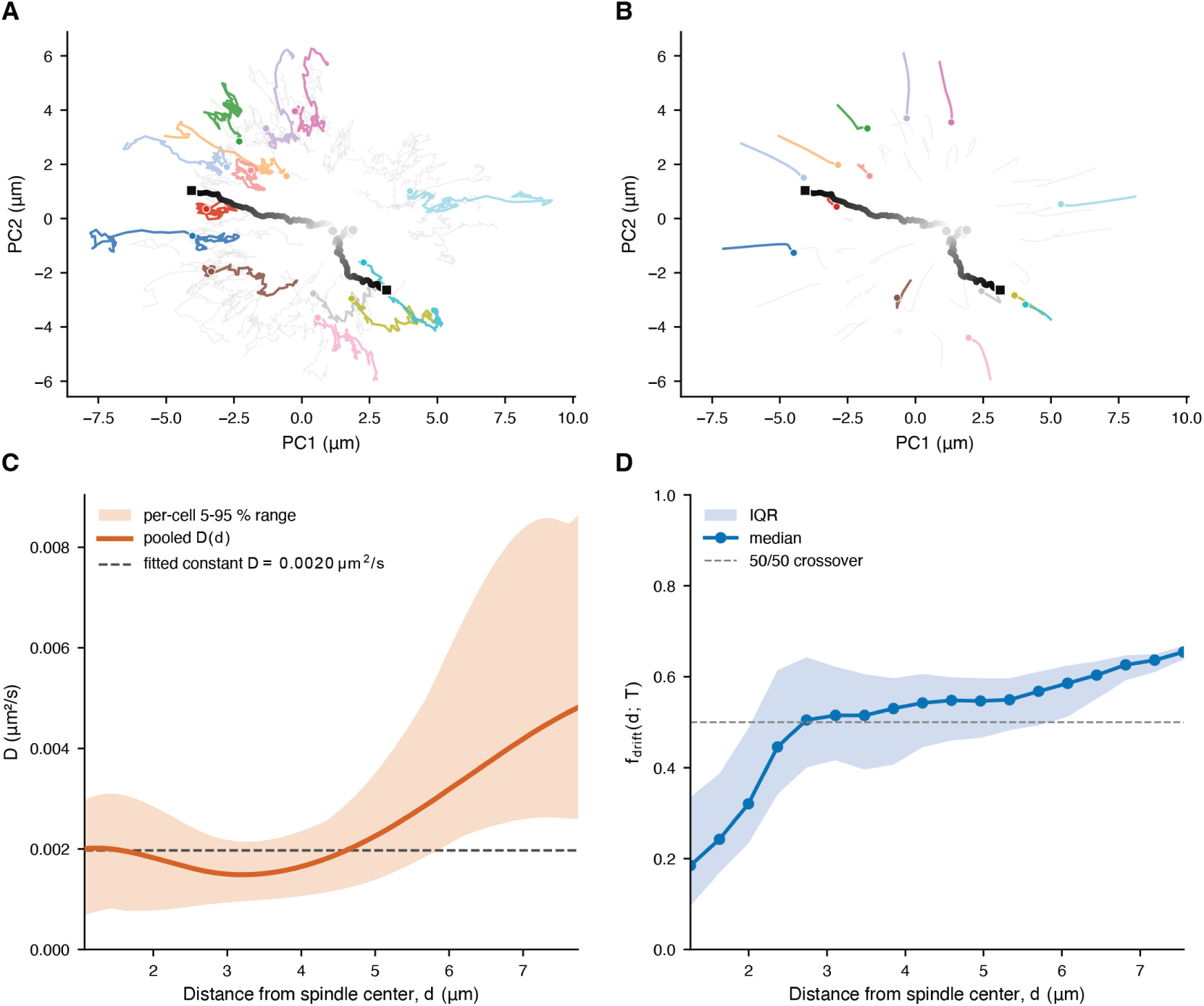
The selected model reproduces chromosome congression. Chromosome motion becomes more drift-dominated with distance from the spindle center. **(A)** Observed centrosome and chromosome (CH) trajectories in a representative control cell (same cell and projection as Fig. 1D). **(B)** Result of a simulation of the selected model (Eq. 1 without the noise term), initialized from the observed CH positions in the first frame and driven by the measured centrosome trajectories. The cell, projection and highlighted CHs are the same as in A. In A and B, centrosomes are shaded from light to dark grey with time (circles, start; squares, end); highlighted CHs are shown with one color each (color does not encode time). **(C)** Effective diffusivity *D d* of CHs as function of three-dimensional distance *d* from the spindle center. Solid line, fit pooled over all cells; band, 5–95% range of per-cell fits; dashed line, best-fit constant diffusivity (*D* = 2 × 10^−3^ *μ*m^2^ s^−1^). *D* increases with distance from the spindle center. **(D)** Drift fraction 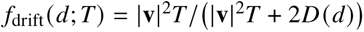, where **v** is the drift velocity predicted by the fitted kernels and *D d* is taken from Fig. 3C. This is the fraction of the mean squared displacement along the drift direction over a time window *T* that is due to deterministic drift rather than diffusion; *T* = 150 s is the duration of the steady elongation phase (Fig. 4A). The dashed line marks *f*_drift_ = 0.5, where the squared drift displacement |**v**|^2^*T* ^2^ equals the diffusive variance 2*D*(*d*)*T*. Values were computed for each chromosome and frame and binned by *d*; solid line, bin median; band, interquartile range. Drift exceeds diffusion beyond *d* ≈ 2.7 *μ*m.

**Figure 4:**
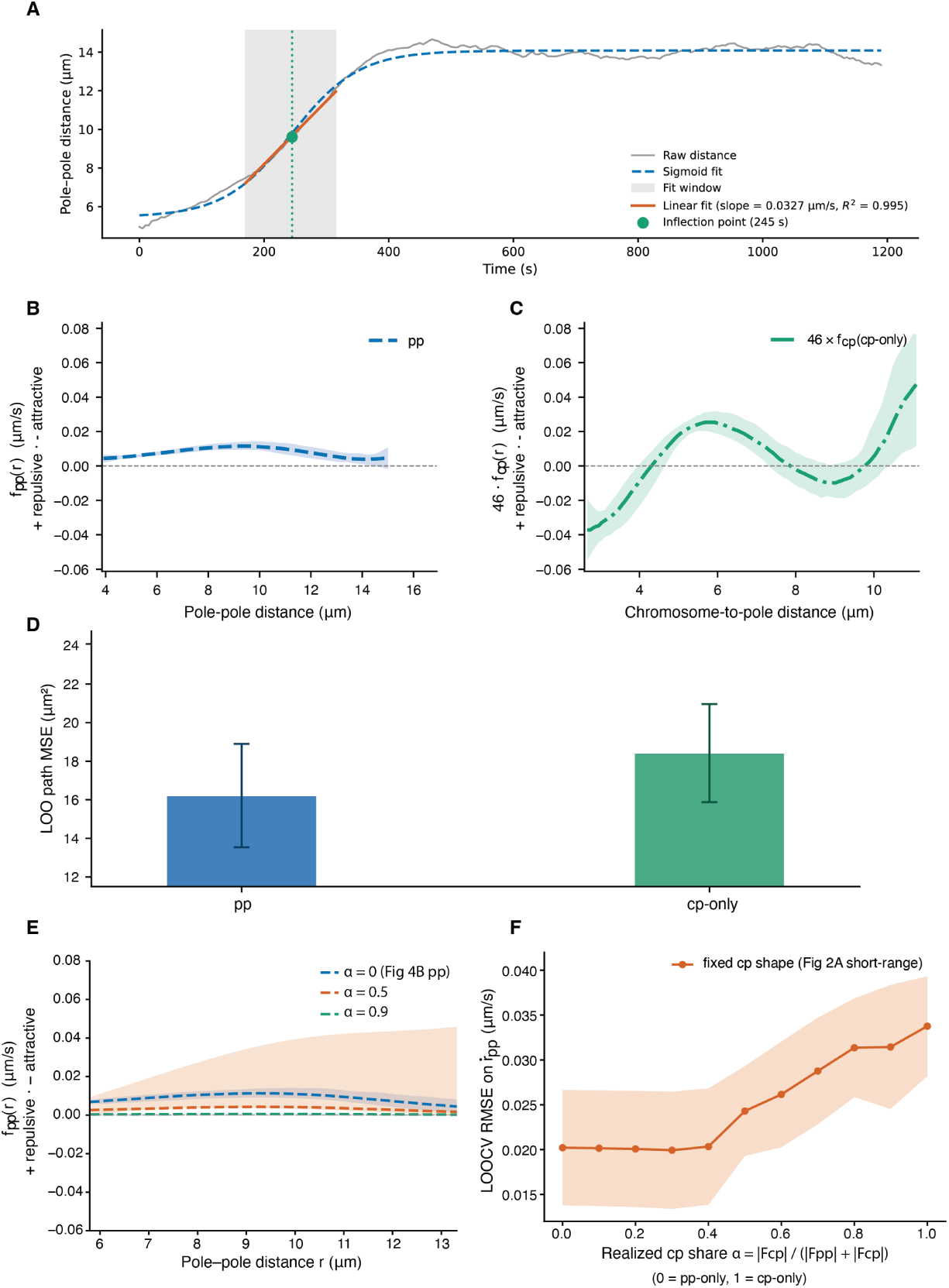
Pole separation is dominated by the interpolar force. **(A)** Pole–pole distance versus time in a representative control cell (grey line; 5 *s* per frame). A four-parameter logistic function (blue dashed line) fit to the full trace locates the steepest phase of pole separation. Its inflection point (green) centers a 150 *s* analysis window (grey shading). Within this window, the distance is well described by a straight line (red), so the poles separate at a nearly constant rate (*R*^2^ ≈ 0.99 for all *n* = 9 control cells). The same window is used for the elongation-rate statistics in Fig. 5B. **(B)** Pole–pole kernel *f*_pp_(*r*) as function of pole–pole distance *r*, inferred from pole velocities with the interpolar term alone (pp model). **(C)** Chromosome–pole kernel inferred with the chromosome–pole term alone (cp-only model), plotted as 46 *f*_cp_(*r*) to place it on the scale of the single interpolar term in Fig. 4B. This scaling is for display only. The actual chromosome contribution to pole velocity is smaller, because chromosomes pull from different directions and their forces partially cancel. **(D)** Leave-one-cell-out path error (mean squared error in 3D pole position given by a simulation of the deterministic model over the same frames as in Fig. 2C) for the pp and cp-only models. Bars, mean over held-out cells; error bars, standard error. **(E)** Interpolar kernel *f*_pp_(*r*) inferred when the chromosome–pole term is included at an indicated fraction *α* of the total force (defined in Fig. 4F). The shape of the chromosome–pole kernel is fixed to the one learned from chromosome motion (Fig. 2A, short-range model), and its amplitude is constrained to be non-negative. *α* = 0 recovers Fig. 4B. As *α* increases, *f*_pp_ decreases in magnitude but remains nearly independent of *r*. Lines, fits to all cells. In B, C and E, kernels are in velocity units; positive values denote repulsion (pole separation), negative values attraction, and the dashed line marks zero. Bands are 5–95% bootstrap confidence intervals. **(F)** Leave-one-cell-out RMSE of the predicted pole-separation rate *r*_pp_ versus the realized chromosome–pole fraction 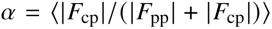 (*α* = 0, interpolar only; *α* = 1, chromosome–pole only). The chromosome–pole kernel shape is fixed as in Fig. 4E. Points, mean over held-out cells; band, ±1 s.d. across folds. The error is nearly flat for *α* ≲ 0.4 and rises for larger fraction of the chromosome-pole forces.

### Pole-separation rate

For each cell, the pole–pole distance was fit by a four-parameter logistic function, 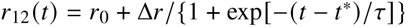, whose inflection point *t*^∗^ marks the fastest separation. The analysis window comprises the 30 frames (150 s) centered on *t*^∗^. Within this window, *r*_12_(*t*) is nearly linear (Fig. 4A). This window is used for the elongation-rate statistics below; the pole models (Fig. 4B–F) were fit on the same frames as the chromosome models. Frame-to-frame elongation velocities, 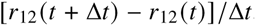, within the window (29 per cell) were pooled by condition to obtain the mean, the s.d. and the signal-to-noise ratio (SNR; mean/s.d.). Variances were compared with the Brown–Forsythe test.

### Stochastic model of the interpolar force balance

Following (16, 35), we model the interpolar MTs as *N* antiparallel overlaps that slide apart at a common velocity *V* (Fig. 5A). Overlap *i* carries *m_i_* kinesin-5 motors and *n_i_* kinesin-14 motors. Kinesin-5 motors follow a linear force–velocity relation with stall force *F*_5,_*_i_* and free velocity *V*_5,_*_i_* (35). Kinesin-14 motors, dragged against their minus-end direction, act as passive dashpots with drag coefficient *z*_14,_*_i_*. The force balance

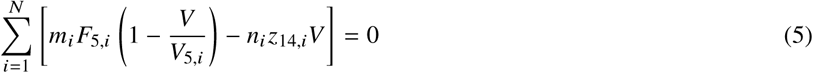

**Figure 5:**
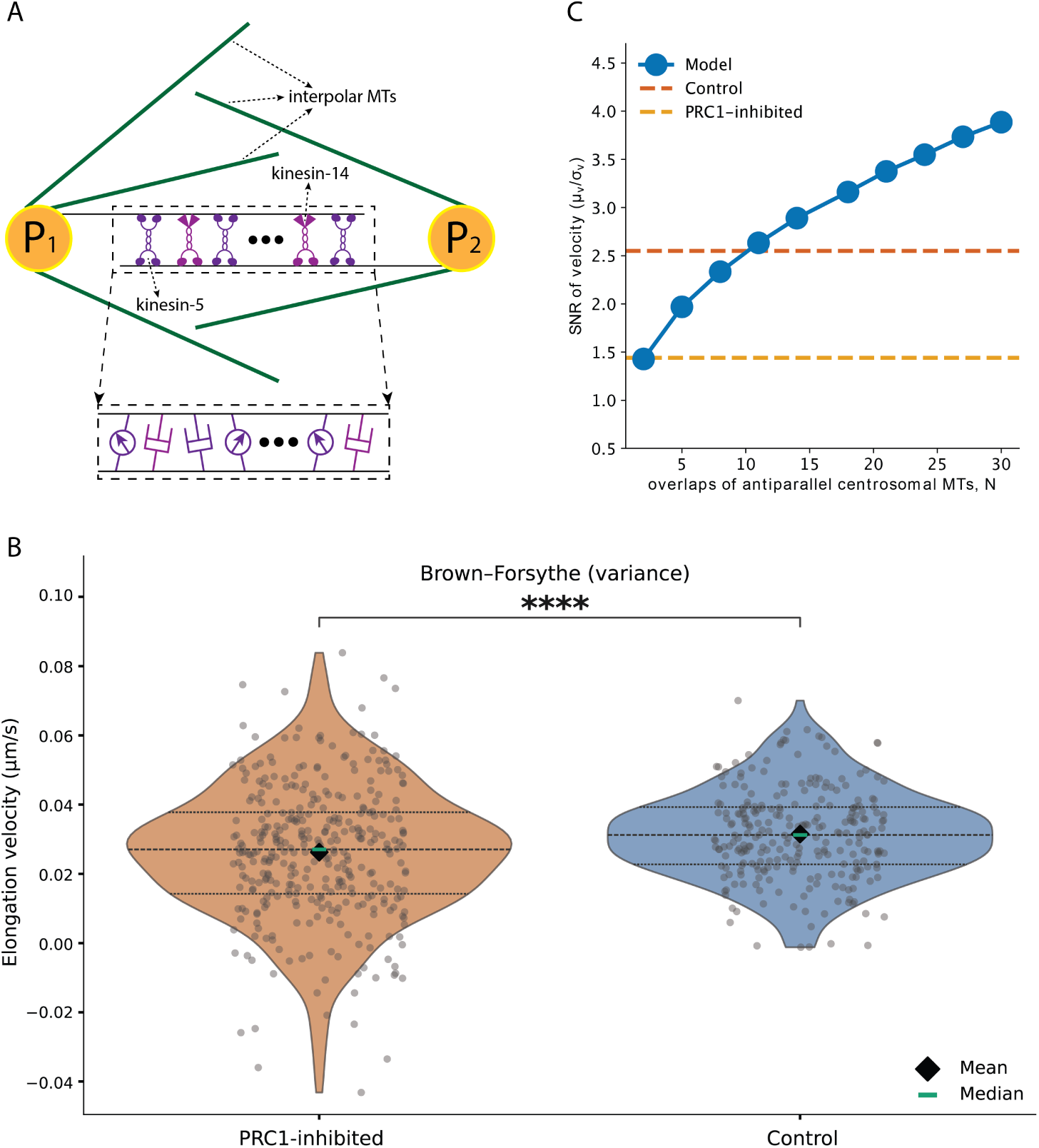
A kinesin-5 / kinesin-14 force balance accounts for the mean and variability of pole separation. **(A)** Model of the interpolar force balance (Eq. 5). The poles (*P*_1_, *P*_2_) are connected by *N* antiparallel interpolar MT overlaps. Each overlap engages on average *m* kinesin-5 and *n* kinesin-14 motors. Inset: mechanical equivalent of one overlap. Each kinesin-5 acts as a force generator with a dashpot (stall force *F*_5_, free velocity *V*_5_). Each kinesin-14, dragged against its minus-end direction, acts as a passive dashpot with drag coefficient *z*_14_. **(B)** Frame-to-frame pole–pole elongation velocities within the analysis window of each cell (Fig. 4A) in PRC1-depleted (vermilion; *n* = 348 velocities from 12 cells) and control (blue; *n* = 261 velocities from 9 cells) spindles. Violins show kernel density estimates with quartiles (dashed lines); grey points, individual velocities; black diamonds, means; green bars, medians. PRC1 depletion increases the standard deviation of the spindle elongation velocity 1.5-fold (0.0195 vs. 0.0131 *μ*m *s*^−1^; Brown–Forsythe test, *p* = 9.7 10^−8^; ****, *p* < 10^−4^). The signal-to-noise ratio (SNR; mean/s.d.) therefore falls from 2.55 in control to 1.44 in PRC1-depleted spindles. **(C)** Model-predicted signal-to-noise ratio (SNR) of the spindle elongation rate versus the number of interpolar MT overlaps *N* (blue), computed from 10^5^ Monte Carlo realizations of Eq. 6 for each *N*. In each realization, the per-overlap motor numbers *m* and *n* are drawn from Poisson distributions with means fixed at their best-fit values, and *F*_5_, *V*_5_ and *z*_14_ are drawn from lognormal distributions with the means and s.d. in SI Table S1 (Methods). Dashed lines, measured SNR in control and PRC1-depleted spindles (Fig. 5B).

and

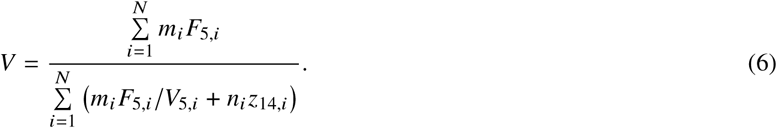

The numerator is the total kinesin-5 stall force. The denominator is the total effective drag, which combines the force–velocity slope of kinesin-5 with the friction of kinesin-14. We identify *V* with the pole-separation rate, neglecting MT poleward flux and the smaller astral, drag and chromosome–pole contributions (see Results).

The per-overlap quantities are independent random variables, identically distributed across overlaps. Motor numbers are Poisson distributed, *m_i_* ∼ Poisson (*λ_m_*) and *n_i_* ∼ Poisson (*λ_n_*). *F*_5,_*_i_*, *V*_5,_*_i_* and *z*_14,_*_i_* are lognormal with means *μ_F_*, *μ_V_*, *μ_z_* and s.d. *σ_F_*, *σ_V_*, *σ_z_*; these are moments of the variables themselves, not of their logarithms. Lognormal distributions keep these quantities positive and right-skewed, as observed in single-molecule measurements (36, 37). We nondimensionalize forces by the mean stall force *F*_0_ and velocities by the mean free velocity *V*_0_, so that *μ_F_* = *μ_V_* = 1 and *σ_F_*, *σ_V_* become coefficients of variation. The SNR, ⟨*V*⟩/s.d.(*V*), is dimensionless and independent of *F*_0_ and *V*_0_. Because *V* is a ratio of sums over *N* independent overlaps, t<u>h</u>e delta method gives, to leading order in 1/*N*, a mean independent of *N* and a variance proportional to 1/*N*, so that SNR 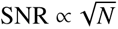 (Supplemental materials).

Parameters were identified by a Monte Carlo search (Supplemental materials). We drew 10^5^ parameter sets uniformly from prescribed ranges and discarded those that violated interval constraints on ⟨*V*⟩. The remaining sets were ranked by the normalized squared deviation of the delta-method SNR from the control value, with *N* fixed at 10. The best-scoring set was *λ_m_* ≈ 30, *λ_n_* ≈ 7.6, *σ_F_* ≈ *σ_V_* ≈ 1.2, *μ_z_* ≈ 0.58 and *σ_z_* ≈ 0.13. The predictions in Fig. 5C and in the SI sensitivity analysis, where one parameter at a time is varied about this baseline, were computed by direct Monte Carlo simulation of Eq. (6) with 10^5^ realizations per parameter value. We used simulation here because the delta method is inaccurate at small *N* and for large parameter spreads.

### Parameter values

For order-of-magnitude force estimates (Results, Discussion) we used the following values. Spindle motors generate forces of 1–10 pN and move at ∼0.1 *μ*m s^−1^ (38). Kinesin-5 has a stall force of ∼1.5 pN (21) and a free velocity of 0.1–0.25 *μ*m s^−1^ (21, 39, 40). Dynein has a stall force of 2–4 pN, a free velocity of 1–2 *μ*m s^−1^ (41), and a nearly linear force–velocity relation (42). The measured spindle elongation rate is ∼0.03 *μ*m s^−1^ (this study).

## RESULTS

### CHs are attracted to spindle poles at long distances and repelled from CSs at short distances, while inter-CH interactions are limited to short-range steric repulsion

We start with learning forces driving CHs movements by using the approach mathematical details of which are explained in Methods (Eq. (1)). In short, from the data, we know positions of all CSs and CHs at many time points, as well as movements of all CSs and CHs from each of these times to the next one. We go over vast number of possible pairwise, distance-dependent forces and find those that approximate the measured movements best. We consider the following possibilities: 1) it is widely accepted that CHs interact, directly or indirectly (5) with long centrosomal MTs, which suggests a CS-CH force (G in Eq. (1), Fig. 1C). 2) It is possible that CHs interact with integrated MT network of the spindle, and so are attracted to (and/or repelled from) the spindle center (midpoint between two poles, force H in Eq. (1), Fig. 1C). 3) It is conceivable that CHs interact with each other through short MTs associated with their kinetochores (43) or through chromatin links (44). Also, mono-oriented CHs were observed to congress to the spindle through lateral attachments to kinetochore fibers of bi-oriented CHs (26). All this raises a possibility of an effective pair-wise CH-CH interaction force (F in Eq. (1), Fig. 1C). In fact, there is the following theoretical possibility: from the beginning, the center of mass of the CH collective is relatively close to the middle between two CSs, so if CHs are effectively attracted to each other, they can congress to the spindle center even without or with a weak interaction with the spindle poles.

We considered four scenarios: 1) each CH interacts only with the spindle poles (*F* ≡ 0, *G* ≠ 0, *H* ≡ 0); 2) each CH interacts only with the spindle center (*F* ≡ 0, *G* ≡ 0, *H* ≠ 0); 3) each CH interacts with the spindle poles AND with each other (*F* ≠ 0, *G* ≠ 0, *H* ≡ 0), without any constraints on the magnitude and distance dependence of the CH-CH force; 4) each CH interacts with the spindle poles AND with each other, BUT with the CH-CH force restricted to be the short-distance repulsion arising simply from the steric repulsion between the hard objects of CH size.

The results are shown in Fig. 2. First, “CHs interact with poles + with each other without constraints” model indeed confirms the intuition that CH-CH attraction can bring the CHs to the spindle when the long-distance CS-CH attraction is almost zero (Fig. 2A). However, this model predicts that CHs attract each other at very large (in the range of ∼10*μ*m) distances (Fig. 2B), which is biologically very hard to justify. Plus, this model has a larger held-out error than two of the other models (Fig. 2C), so we reject it. Second, “CHs interact with the spindle center model” makes sense in principle: its analysis shows that the CHs are attracted to the spindle center at medium distances and do not interact with the center at small and large distances (Fig. 2A). However, this model has the largest held-out error of the four (Fig. 2C), and we reject this model. Third, the model “CHs interact with poles only” is statistically reasonable (Fig. 2C) and produces a clear prediction: the CS-CH interaction is a strong repulsion at small distances and weak and almost constant (barely distance dependent) attraction at greater distances (Fig. 2A). Finally, adding hardcore CH-CH repulsion to the CS-CH interaction reduces the error even more (Fig. 2C), without changing the predicted CS-CH interaction (Fig. 2A). Thus, we accept that last model.

To illustrate that this model works, we use the measured pole trajectories to calculate several CH trajectories (starting from the experimentally measured CHs’ initial positions) by using Eq. (1) with the learned parameters and functions (Fig. 3B), and then compare the computed CH trajectories with the experimentally measured ones (Fig. 3A). They agree qualitatively and even semi-quantitatively. The result demonstrates the predictive power of the learned model.

### Random movements of the CHs are faster farther away from the spindle

From fitting the model (Eq. (1)) to the data, we also were able to estimate the magnitude of random, diffusion-like CH movements (Fig. 3C). We found that these random movements account for displacements of tenths of a micron over the characteristic 100-seconds time interval during early prometaphase. Interestingly, these movements are faster farther away from the spindle (Fig. 3C), which is in agreement with the reported importance of random CH movements for the CH congression to the spindle (6). Note that if we use the learned CS-CH force to compute the average, distance-dependent, drift to (at long distances) and away from (at short distances) the spindle, and estimate the ratio of the deterministic drift and random movements, then the net CH movement is more deterministic and centripetal in nature farther away from the spindle and more random in and near the spindle (Fig. 3D), despite the increase of the randomness with the distance. Thus, the average centripetal congression is an increasing function of the distance overwhelming the increase of the randomness with the distance.

### The interpolar forces dominate the net CS-CH force

Now that we established the model for the CS-CH force, we use the data to analyze the poles’ movements, using the same general idea as above: knowing positions and incremental displacements of all spindle elements, we can find the most likely forces by minimizing approximation errors in the model given by Eq. (2) in Methods. In this model, the geometric sum of three forces drives the poles: 1) the net interpolar force (P, Fig. 1C), 2) the net astral force (C, Fig. 1C), 3) the net force from all CHs (G, Fig. 1C) – according to the third Newton’s law, the CS-CH force applied to each CH is accompanied by the opposite force on respective pole. The sum of all these forces is balanced by 4) the effective drag on the pole movement from the intracellular environment (Eq. (2)) (25).

Of these four forces, it is hard to discriminate against the passive poles’ drag and the astral force using only the analysis of the movements. However, the following two arguments make the force analysis simpler. We measured the rate of the spindle length increase and found it to be remarkably constant over the time interval longer than 100 seconds (Fig. 4A). It would be hard to imagine how the net astral force and drag could be a significant part of the total force balance and, at the same time, constant over such a long time, during which the poles travel through very different parts of the cell (Fig. 1B). During these pole excursions, stochastic interactions of astral MTs with the cell cortex are supposed to sum up to a significantly varying in time, not constant force (11, 45–47). Similarly, it would be hard to imagine these two different forces (net astral force and drag) cancel each other variations. This argument suggests that these two forces are much smaller than the principal interpolar forces (see below). Indeed, the net astral and drag forces are not likely to exceed ∼100 pN (see the Introduction and Discussion), while the principal interpolar forces are likely to be several hundreds to thousands of pN ((16) and Discussion). Thus, we assume that the net astral (C in Eq. (2)) and drag forces can be omitted.

We could use similar arguments against a significant net CS-CH force because such net force, ∼100 pN (see the Discussion), is also supposed to be an order of magnitude smaller than the principal interpolar forces, but we prefer to make a more rigorous argument, because unlike the pole’s drag force and the astral forces, which we cannot learn from the data, we already learned the distance dependence of the CS-CH force (Fig. 2).

Therefore, we first examined two models: 1) the whole force comes from the pole-pole interaction; the net CS-CH force is negligible (*P* ≠ 0, *G* = 0); 2) the whole force comes from the net CS-CH force; the pole-pole interaction is negligible (*P* = 0, *G* ≠ 0). Fits of these models to the data are shown in Fig. 4B,C. Fig. 4B shows that expected, not very sensitive to distance, net interpolar force alone could explain the spindle elongation. Fig. 4C, on the other hand, shows that the distance dependence of the CS-CH force would show very strange nonmonotonic behavior that is most likely unbiological. Moreover, the first model produces a smaller error of approximation (Fig. 4D). Thus, the net interpolar force is definitely a significant factor in the spindle elongation.

Next, to evaluate if the net CS-CH force could be a non-negligible part of the sum of the interpolar and CS-CH forces, we considered a third model, in which a percentage (1 − *α*) gives the amplitude of the net interpolar force, and percentage *α* gives the amplitude of the net CS-CH force, and importantly, the distance dependence of the individual CS-CH force is as learned from the data (Fig. 2A). We found that the insensitivity of the interpolar force to the distance is preserved as the fraction of it varies (Fig. 4E), and the interpolar repulsion becomes weaker, as the contribution from the CHs increases. This means that the net CS-CH force repels the poles from each other, which makes sense, considering that most of CHs are between the poles and that the repulsive part of the CS-CH interaction is greater than the attractive part (Fig. 2A).

Most importantly, the error of approximating the data starts to increase as the fraction of the net CS-CH force exceeds 40% (Fig. 4F), supporting the argument made above that the net CS-CH force is much smaller than the interpolar forces. The data do not allow to deduce the exact percentage of the net CS-CH force in the total balance, but we use the semi-quantitative arguments and analysis here to assume that this force is dominated by the interpolar forces.

### Tug-of-war between Kinesin-5 and Kinesin-14, with tens of kinesin motors at the tens of antiparallel MT-MT overlaps on the spindle equator can explain the pole-pole repulsion

We used measurements of the spindle elongation to measure the rate of the pole-pole segregation and its fluctuations (Fig 4A) during the long-time interval (more than 100 seconds) in early prometaphase around the time of the fastest segregation. Here, we use a simple model inspired by the first quantitative analysis of prometaphase (16) and of Kinesin-5 / Kinesin-14 force balance (35) to explain these measurements.

We assume that PRC1 crosslinkers maintain *N* overlaps of antiparallel MTs at the spindle equator. Following (16), we posit that at each of these *N* overlaps, *m* Kinesin-5 motors, on average, and *n* Kinesin-14 motors, on average, are engaged (Fig. 5A). Because the spindle elongates, Kinesin-5 motors sliding the antiparallel interpolar MTs apart are winning. It was shown in (35) that in this regime, Kinesin-5 motors are characterized by the linear force velocity relation, such that when the motors are stalled, they collectively generate a force *mF*_5_ (*F*_5_ is per-motor stall force), and when they move freely against zero force, they slide the poles apart with velocity *V*_5_. On the other hand, Kinesin-14 motors, which are dragged against their normal minus-end direction, act as passive viscous dashpots, each adding effective viscous drag *z*_14_*V* proportional to the spindle elongation rate *V*. The resulting force balance, taking into account both kinesins’ forces, neglecting the smaller drag, astral and CS-CH forces, is expressed by Eq. 5 (Methods).

In the model, we treat model number parameters *n*, *m* and motor mechanical parameters *F*_5_, *V*_5_, *z*_14_ as random numbers. More specifically, in the calculations and simulations, parameters *F*_5_, *V*_5_, *z*_14_ are drawn from lognormal distributions (Methods), while parameters *n*, *m* are treated as the result of Poisson processes that stem from the assumed MT and motors’ dynamics (Methods). We vary the averages of parameters *n*, *m* and solve Eq. 5 analytically and numerically to investigate which parameters produce the observed fluctuations of the spindle elongation rate (Fig. 5B). With *N* fixed at 10 (ten interpolar MT bundles), we find that *m* ≈ 30 Kinesin-5 and *n* ≈ 8 Kinesin-14 motors per bundle fit the data best (Supplemental materials).

Translated into forces, this result means that several hundreds of Kinesin-5 motors are mainly opposed by several hundreds of Kinesin-14 motors, and this opposition in the range of 500 pN from two sides almost cancel each other, setting the observed rate of spindle elongation. The additional ∼100 pN of astral force minus ∼100 pN of drag plus a few tens pN of the net CS-CH force perturb the dominant Kinesin-5/Kinesin-14 balance and poles’ segregation rate by 20% at most (Methods).

To test this prediction, we perturbed Protein Regulator of Cytokinesis 1 (PRC1) – a vital crosslinker that binds and bundles interpolar MTs stabilizing the central zone of the mitotic spindle (17). We measured the average rate of spindle elongation and its variance in PRC1-depleted spindles (Fig. 5B). We found that the average rate of pole-pole separation remains the same in the perturbed cells, while the level of noise of the spindle elongation rate increases. This perturbation does not affect mechanical parameters of kinesin motors and likely does not affect parameters *n* and *m* (as those depend on interpolar MT lengths, not on their numbers). Most likely, PRC1 depletion decreases the number of interpolar MT bundles, *N*. (In principle, this inhibition could also decrease the passive drag of sliding interpolar MTs apart, but this is less likely to cause a significant effect because of abundance of other MT associated proteins). We examined dependence of the model’s behavior on parameter *N*, and indeed found that as *N* decreases, the average spindle elongation rate stays the same, while its variance increases (Fig. 5C), supporting the model.

## DISCUSSION

Understanding mitotic spindle was achieved, in part, by assuming mechanical interactions between the spindle parts, simulating the resulting dynamics and comparing them to the data (48–51). As more detailed data become available, a complementary modeling approach – using data on trajectories of CHs and CSs to constrain a general model – started to work (52, 53). Here, we started with a data-based, and finished with the traditional approach to understand force balances in the mitotic spindle that are not well established. Our analysis suggests that, in early prometaphase, the CHs do not interact nontrivially with each other, other than through the short-range steric repulsion.

Our main finding is that the CHs are attracted to each of the CSs at CS-CH distances greater than ∼4 *μ*m. The effective attraction force is almost constant, insensitive to the CS-CH distance, at CS-CH distances greater than ∼6 *μ*m. At the short range, below ∼4 *μ*m, CHs are effectively repelled from the CSs. In addition, CHs undergo random diffusive movements that are more vigorous farther away from the spindle. Molecular interpretation of this result is straightforward: minus end-directed dynein motors drive CHs poleward at greater distances, while kinesin- and/or polymerization-powered anti-poleward forces dominate near the poles (54). The learned distance dependence of the CS-CH force seems to be optimal for stabilizing the CHs on the spindle equator.

By the third Newton’s Law, this net CS-CH force is also applied to both CSs. Our analysis shows that this net CS-CH force effectively could repel the spindle poles from each other because many CHs are relatively close to the CSs in early prometaphase. However, according to the model, the data suggest that this is a minor (less than 40%) part of the pole segregation forces, while the principal forces come from the interpolar MT motors, specifically, by hundreds of Kinesin-5 motors opposing hundreds of Kinesin-14 motors distributed over ten interpolar MT bundles with tens of kinesin motors per bundle. We note that effective viscous drags from MT crosslinkers (55, 56) and dyneins (57) on the interpolar MT overlaps could add to the Kinesin-14 action. We argue that the highly variable astral MT-dynein force is relatively small and changes the interpolar force balance little. The noise analysis suggests that hundreds of kinesin-5 and kinesin-14 motors generate the net repulsive interpolar force, meaning that kinesin-5 overwhelms kinesin-14.

Our analysis also suggests that the interpolar force balance in early prometaphase depends little on the distance between the poles. This means that the pole-pole segregation is robust, largely insensitive to regional variations inside the cell and to complex process of CH congression, which underscores the importance of the pole-pole segregation for the spindle assembly (5, 6). On the first glance, this is surprising considering the dynamic spindle architecture, but the explanation is rather simple: the average rate of pole-pole segregation is determined mostly by force-velocity relations of the opposing kinesin motors and ratio of the opposing motor numbers, rather than the total motor numbers dependent on the exact spindle structure. The PRC1 depletion result underscores this point: PRC1 depletion decreases crosslinking of the antiparallel interpolar MTs, which effectively diminishes the total numbers of the opposing kinesin motors on the antiparallel interpolar MTs, but the average spindle elongation rate does not change in a statistically significant way.

To conclude, our analysis posits the following hierarchy of the spindle forces in early prometaphase. The spindle poles are segregated by hundreds of opposing kinesin motors generating ∼500 pN of opposing forces. Balancing these large forces results in a steady segregation of the poles at an almost constant rate not very sensitive to the spindle length. An order of magnitude weaker forces from the poles, meanwhile, push and pull the CHs individually resulting in the CH congression to the spindle. The CH movements are ‘slaved’ to the poles and have little effect on the spindle elongation. The spindle is rotated and its center-of-mass is positioned by astral forces that are weaker than the opposing interpolar forces.

These findings agree with conclusions on principal motor balances made in (16). The distance dependence of the CS-CH force agrees with the analysis of the CH congression in our previous study (6). Our prediction of the order of ten of interpolar MT bundles seems to agree with discussion of data in (19).

There are also thought-provoking differences with a couple of previous studies. Thus, one early estimate of the total number of kinesin motors leans to thousands (27), not hundreds, as in our result, but this early estimate was based on the maximal possible number of motors from geometric point of view, rather than from the analysis of mechanics. Further, Drosophila embryos are known to contain large stores of pre-produced protein to support the extremely rapid divisions of numerous spindles within the syncytium. Our conclusions are also not consistent with the reported lack of correlation in the movements of centrosomes during anaphase in cultured newt lung cells (58). This observation suggests that centrosomes are displaced by dynein-mediated pulls along the astral MTs rather than by the pushing forces in the spindle midzone. However, the lack of pushing forces in this experimental system likely reflects the unique architecture of the spindle in newt cells where the distance between the spindle poles may exceed 50 *μ*m which severely decreases MT overlap near the spindle equator. Indeed, the authors of (58) conceded that their observations in newt cells contradict findings from different experimental systems.

The models we posit in this study have several limitations. First, they are limited to early prometaphase – as different kinds of motors switch on and off between the mitotic phases (24), it would be futile to extend the exact same model to explain, for example, the transitions from prophase and to metaphase when the spindle starts and stops elongating, respectively. However, methods similar to those developed in this study could guide understanding of other dynamic mitotic stages, such as anaphase (52). Besides, certain modeling assumptions will have to be examined further. Effective CS-CH forces, for example, may be not pairwise and not reducible to simple distance-dependent forces if detailed MT dynamics and MT-motor interactions are taken into account. More mechanically complex load-dependent motor detachments (16, 59) and/or depolymerization of MTs at the poles in response to forces (60) could alter the force balance equations. The fact that some CHs become integrated into the spindle much faster than other CHs could significantly affect spindle MT networks and force balances (4). Processes other than MT-based motors pushing and pulling (7), as well as interactions with cytoskeletal and structural elements of the spindle other than MTs (16), could also contribute to the mechanical equilibrium in the spindle.

Last, but not least, it remains to be determined what are the force balances in prometaphase when spindles are organized under different design principles (61, 62). It would also be imperative to examine scaling laws of prometaphase spindles in different organisms, similar to how this was done for metaphase spindles (63, 64). Ultimately, direct force measurements together with detailed microscopy will be needed to test the mechanics in prometaphase.

## AUTHOR CONTRIBUTIONS

W.S., C.M., A.K. and A.M. designed the research. W.S., C.M., F.R. and A.K. analyzed the data. W.S. and C.M. performed the research. W.S., C.M. and A.M. wrote the paper.

## ACKNOWLEDGMENTS

This work was supported by the NIH NIGMS grant GM130298 (A.K.), NSF Grants DMS 1953430 (A.M.) and CAREER DMS-2545859 (C.E.M.) and NSF/NIGMS Grant DMS-2451263 (C.E.M.).

## SUPPLEMENTARY MATERIAL

Supplementary material is provided as a separate file accompanying this preprint.

