## Supplemental Materials for "Hierarchy of spindle forces in prometaphase"

#### S1 Pole separation in control and PRC1-inhibited cells

Figures S1 and S2 show the pole–pole distance $r_{12}(t)$ in every control ($n = 9$) and PRC1-inhibited ($n = 12$) cell analyzed in the main text. For each cell, the full trace was fit by a four-parameter logistic function,

$$r_{12}(t) = r_0 + \frac{\Delta r}{1 + \exp[-(t - t^*)/\tau]}, \tag{S1}$$

where $r_0$ is the initial distance, $\Delta r$ the total increase, $t^*$ the inflection time and $\tau$ the time scale of separation. The separation rate $\mathrm{d}r_{12}/\mathrm{d}t$ is largest at $t^*$. The analysis window comprises the 30 frames (150 s) centered on $t^*$. Anchoring the window to $t^*$, rather than to a fixed time after nuclear envelope breakdown, selects the phase of fastest separation in each cell despite cell-to-cell differences in timing. Within the window, $r_{12}(t)$ is close to linear; the slope and $R^2$ of a linear fit are given in each panel. The same procedure was applied to both conditions. Frame-to-frame elongation velocities, $[r_{12}(t + \Delta t) - r_{12}(t)]/\Delta t$ with $\Delta t = 5$ s, were computed within each window (29 per cell) and pooled by condition (261 control and 348 PRC1-inhibited velocities). Their distributions are shown in main-text Fig. 5B. The ratio of their mean to their s.d. gives the measured SNR (signal to noise ratio) used in Section S2.

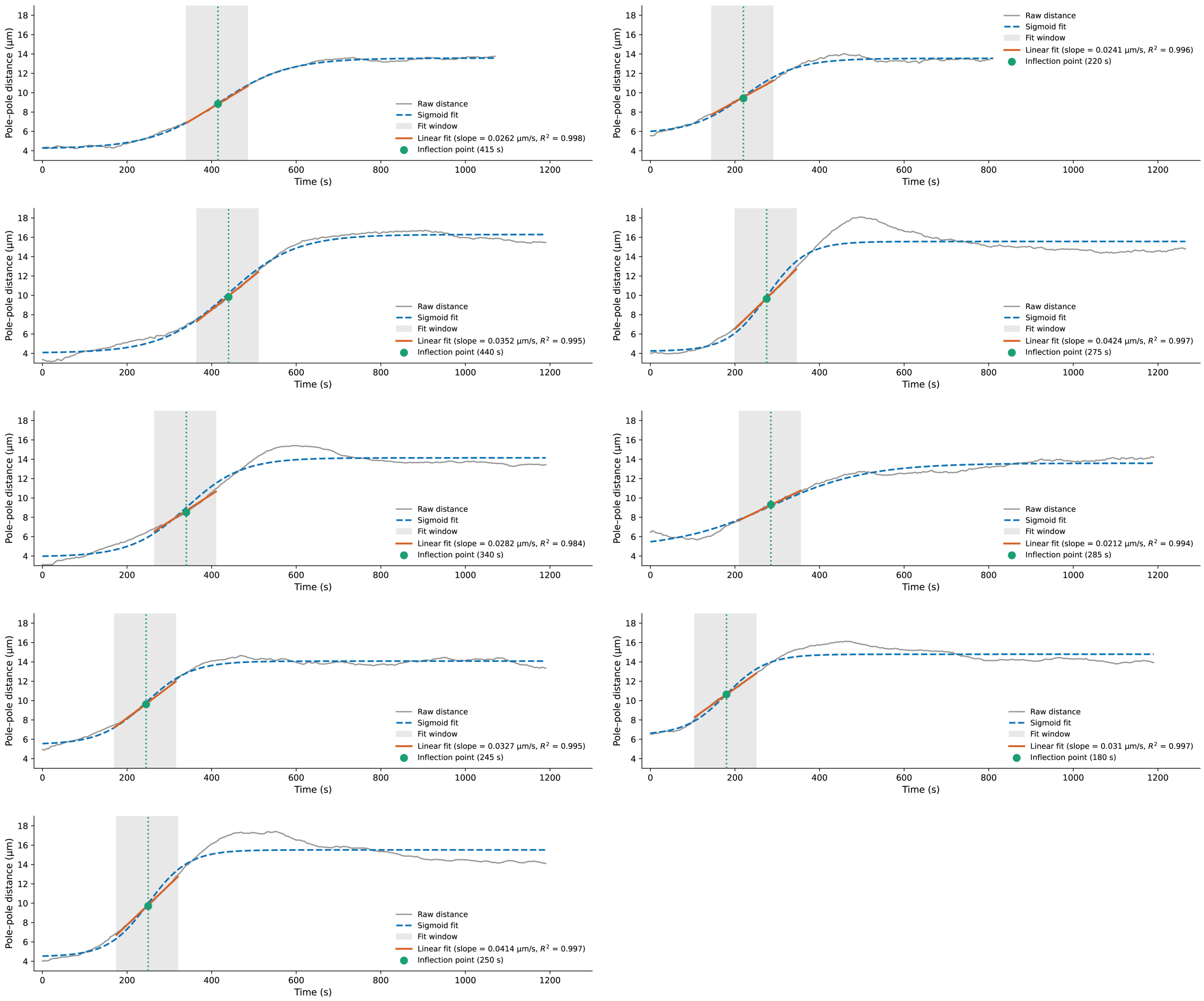


**Figure S1. Pole–pole distance in all control cells.** Each panel shows one of the $n = 9$ control cells. Grey line, pole–pole distance $r_{12}(t)$ (5 s per frame); blue dashed line, four-parameter logistic fit to the full trace (Eq. S1); green dotted line, its inflection point $t^*$; grey shading, the 30-frame (150 s) analysis window centered on $t^*$; red line, linear fit within the window, with slope and $R^2$ given in each panel. Window selection and fitting are described in Section S1.

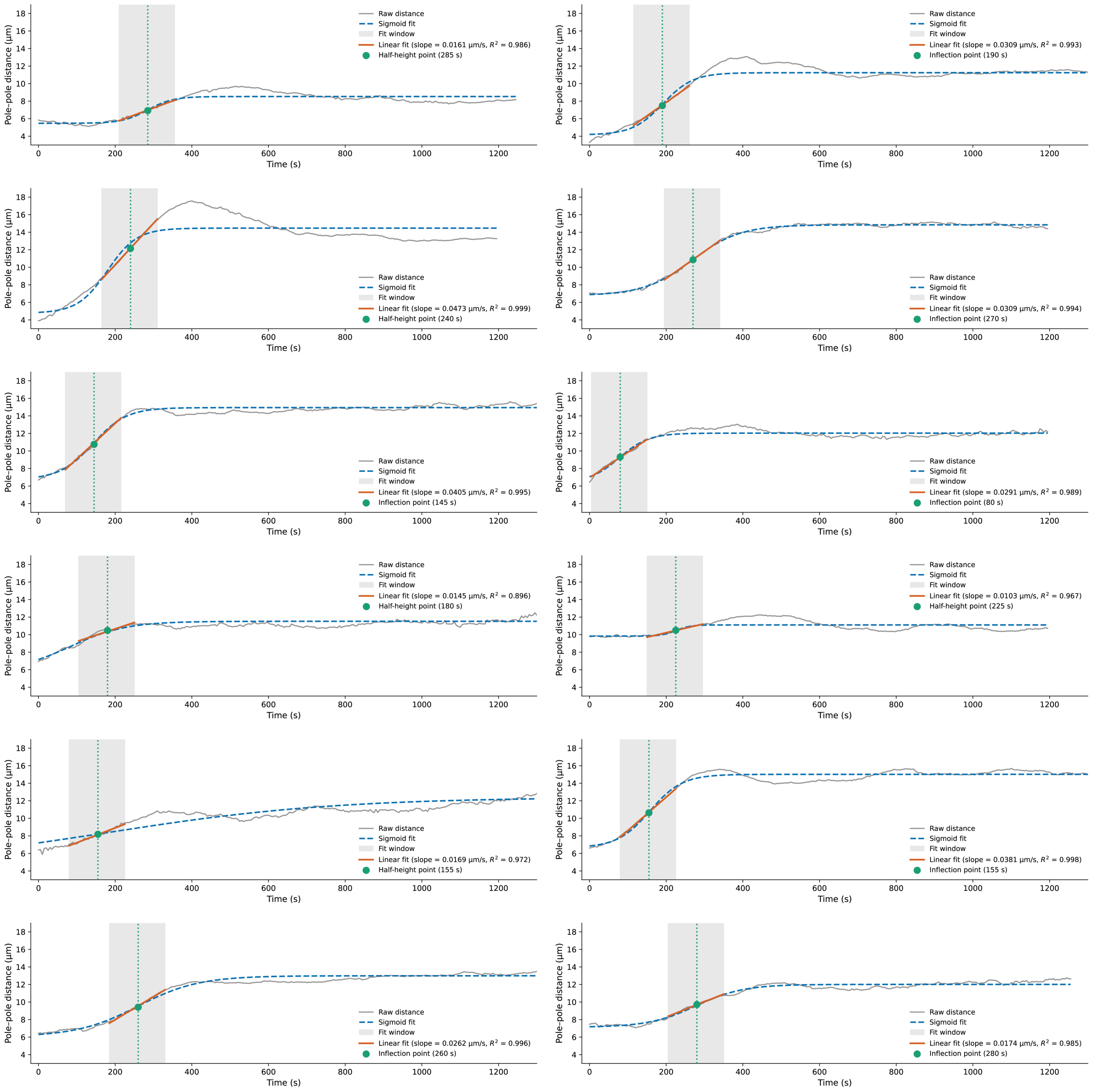


**Figure S2. Pole–pole distance in all PRC1-inhibited cells.** Each panel shows one of the $n = 12$ PRC1-inhibited cells. Grey line, pole–pole distance $r_{12}(t)$ (5 s per frame); blue dashed line, four-parameter logistic fit to the full trace (Eq. S1); green dotted line, its inflection point $t^*$; grey shading, the 30-frame (150 s) analysis window centered on $t^*$; red line, linear fit within the window, with slope and $R^2$ given in each panel. Window selection and fitting are described in Section S1.

### S2 Stochastic model of the interpolar force balance

#### S2.1 Model

The interpolar MTs are modeled as $N$ antiparallel overlaps that slide apart at a common velocity $V$. Overlap $i$ ($i = 1, \ldots, N$) carries $m_i$ kinesin-5 and $n_i$ kinesin-14 motors. The kinesin-5 motors of overlap $i$ share a stall force $F_{5,i}$ and a free velocity $V_{5,i}$ and follow a linear force–velocity relation. The kinesin-14 motors, dragged against their minus-end direction, act as passive dashpots with drag coefficient $z_{14,i}$. The force balance

$$\sum_{i=1}^{N} \left[ m_i F_{5,i} \left( 1 - \frac{V}{V_{5,i}} \right) - n_i z_{14,i} V \right] = 0 \tag{S2}$$

gives

$$V = \frac{X}{Y}, \qquad X = \sum_{i=1}^{N} X_i, \quad X_i = m_i F_{5,i}, \qquad Y = \sum_{i=1}^{N} Y_i, \quad Y_i = \frac{m_i F_{5,i}}{V_{5,i}} + n_i z_{14,i}. \tag{S3}$$

$X$ is the total kinesin-5 stall force. $Y$ is the total effective drag, which combines the slope of the kinesin-5 force–velocity relation with the kinesin-14 friction.

The per-overlap variables are independent and identically distributed across overlaps. Motor numbers are Poisson distributed, $m_i \sim \text{Poisson}(\lambda_m)$ and $n_i \sim \text{Poisson}(\lambda_n)$. $F_{5,i}$, $V_{5,i}$ and $z_{14,i}$ are lognormal, with means $\mu_F$, $\mu_V$, $\mu_z$ and standard deviations $\sigma_F$, $\sigma_V$, $\sigma_z$. Forces are measured in units of the mean stall force $F_0$ and velocities in units of the mean free velocity $V_0$, so that $\mu_F = \mu_V = 1$.

Throughout, $\langle\cdot\rangle$, $\text{Var}(\cdot)$ and $\text{Cov}(\cdot,\cdot)$ denote the mean, variance and covariance over realizations of these random variables. The signal-to-noise ratio (SNR) of the sliding velocity is

$$\text{SNR} = \frac{\langle V \rangle}{\sqrt{\text{Var}(V)}}, \tag{S4}$$

which is dimensionless and independent of $F_0$ and $V_0$. These quantities were computed approximately with the delta method and verified by Monte Carlo simulation (Section S2.2).

#### S2.2 Moments of the sliding velocity

The moments of a single overlap follow from the independence of the variables, the Poisson identities $\langle m_i \rangle = \text{Var}(m_i) = \lambda_m$ and $\langle n_i \rangle = \text{Var}(n_i) = \lambda_n$, and a second-order Taylor expansion of $V_{5,i}^{-1}$ whose mean and variance are approximated by $\mu_W$ and $\sigma_W^2$:

$$\begin{aligned} \mu_W &= \frac{1}{\mu_V} + \frac{\sigma_V^2}{\mu_V^3}, & \sigma_W^2 &= \frac{\sigma_V^2}{\mu_V^4}, \\ \langle X_i \rangle &= \lambda_m \mu_F, & \text{Var}(X_i) &= \lambda_m \mu_F^2 + (\lambda_m^2 + \lambda_m)\,\sigma_F^2, \\ \langle Y_i \rangle &= \lambda_m \mu_F \mu_W + \lambda_n \mu_z, & \text{Cov}(X_i, Y_i) &= \mu_W \, \text{Var}(X_i), \end{aligned} \tag{S5}$$

$$\text{Var}(Y_i) = \text{Var}(X_i)\,\mu_W^2 + \langle X_i \rangle^2 \sigma_W^2 + \text{Var}(X_i)\,\sigma_W^2 + \lambda_n \mu_z^2 + (\lambda_n^2 + \lambda_n)\,\sigma_z^2. \tag{S6}$$

Applying the delta method to $V = X/Y$ and keeping the leading order in $1/N$ gives

$$\langle V \rangle \approx \frac{\langle X_i \rangle}{\langle Y_i \rangle}, \qquad \text{Var}(V) \approx \frac{1}{N} \left[ \frac{\text{Var}(X_i)}{\langle Y_i \rangle^2} + \frac{\langle X_i \rangle^2 \, \text{Var}(Y_i)}{\langle Y_i \rangle^4} - \frac{2 \langle X_i \rangle \, \text{Cov}(X_i, Y_i)}{\langle Y_i \rangle^3} \right]. \tag{S7}$$

Thus $\langle V \rangle$ is independent of $N$ and SNR $\propto \sqrt{N}$.

With $N$ fixed at 10, we identified the parameter values in Table S1, which reproduce the measured control SNR of 2.55. The delta method assumes small relative fluctuations and is inaccurate for the large spreads in Table S1. We therefore verified the parameter values by Monte Carlo simulation. We drew $10^5$ independent realizations of $\{m_i, n_i, F_{5,i}, V_{5,i}, z_{14,i}\}_{i=1}^{N}$, computed $V$ from Eq. (S3) for each, and estimated $\langle V \rangle$ and $\mathrm{Var}(V)$ by the sample mean and variance. This gives $\langle V \rangle \approx 0.46$, $\sqrt{\mathrm{Var}(V)} \approx 0.18$ and SNR $\approx 2.54$, matching the control value. All model predictions reported (main-text Fig. 5C and Fig. S3) were computed by the same Monte Carlo procedure.

**Table S1. Parameter values of the stochastic interpolar model.** Forces are in units of $F_0$ and velocities in units of $V_0$.

| Parameter | Meaning | Value |
|---|---|---|
| $N$ | number of interpolar overlaps | 10 |
| $\lambda_m$ | mean number of kinesin-5 per overlap | 29.965 |
| $\lambda_n$ | mean number of kinesin-14 per overlap | 7.582 |
| $\mu_F$, $\mu_V$ | mean kinesin-5 stall force, free velocity | 1 |
| $\sigma_F$ | s.d. of kinesin-5 stall force | 1.190 |
| $\sigma_V$ | s.d. of kinesin-5 free velocity | 1.186 |
| $\mu_z$ | mean kinesin-14 drag coefficient | 0.583 |
| $\sigma_z$ | s.d. of kinesin-14 drag coefficient | 0.125 |

##### S2.3 Sensitivity analysis

The parameter values in Table S1 are uncertain. Single-motor properties are measured in vitro, the motor numbers per overlap are not measured directly, and motor expression and spindle architecture vary from cell to cell. We therefore tested how robust the model predictions are to changes in each parameter, and which parameters control the mean and the variability of the sliding velocity.

We varied one parameter at a time while holding all others fixed at their baseline values (Table S1): $N$ over integer values from 2 to 30, and each of $\lambda_m$, $\lambda_n$, $\sigma_F$, $\sigma_V$, $\mu_z$ and $\sigma_z$ over 10 evenly spaced values from 0.5 to 1.5 times its baseline. For each value, the mean, standard deviation and SNR of $V$ were computed from $10^5$ Monte Carlo realizations (Fig. S3).

The number of overlaps $N$ controls the variability of $V$ much more than its mean. Between $N = 2$ and $N = 30$, the standard deviation decreases about fourfold, from $\approx 0.42$ to $\approx 0.11$, as expected from $\mathrm{Var}(V) \propto 1/N$ (Eq. (S7)), and the SNR increases from $\approx 1.4$ to $\approx 3.9$. Over the same range, the mean decreases only from $\approx 0.59$ to $\approx 0.42$. Reducing $N$ from its baseline value of 10 to 2 thus raises the mean moderately (by $\approx 30\%$), but more than doubles the standard deviation and lowers the SNR from $\approx 2.5$ to $\approx 1.4$, close to the value measured in PRC1-inhibited spindles (1.44). This agrees with the main-text observation that PRC1 inhibition, which is expected to reduce the number of interpolar MT bundles, increases the noise of spindle elongation much more than it changes its average rate.

Increasing $\sigma_V$ from 0.59 to 1.78 lowers the mean from $\approx 0.69$ to $\approx 0.31$. Each kinesin-5 resists sliding with an effective drag $F_5/V_5$ (Eq. (S3)), which is inversely proportional to its free velocity. Spreading the free velocities at a fixed mean therefore raises the average of $V_5^{-1}$, $\langle V_5^{-1} \rangle = (1 + \sigma_V^2/\mu_V^2)\mu_V^{-1}$ (Eq. (S5)), because the extra drag from motors slower than the mean outweighs the reduced drag from motors faster than the mean. Over this range, $\langle V_5^{-1} \rangle$ increases from $\approx 1.4$ to $\approx 4.2$. The total drag $Y$ increases while the total stall force $X$ is unchanged, so the sliding velocity $V = X/Y$ decreases. The standard deviation changes little, so the SNR falls from $\approx 4.5$ to $\approx 1.9$. A change in $\sigma_V$ therefore lowers the SNR only together with a large drop in the mean, unlike the effect of PRC1 inhibition.

All other parameters have moderate effects. Over their ranges, $\lambda_m$, $\lambda_n$, $\sigma_F$, $\mu_z$ and $\sigma_z$ change the mean by less than 10% (0.42–0.48) and keep the SNR between 2.3 and 2.9, close to the control value.

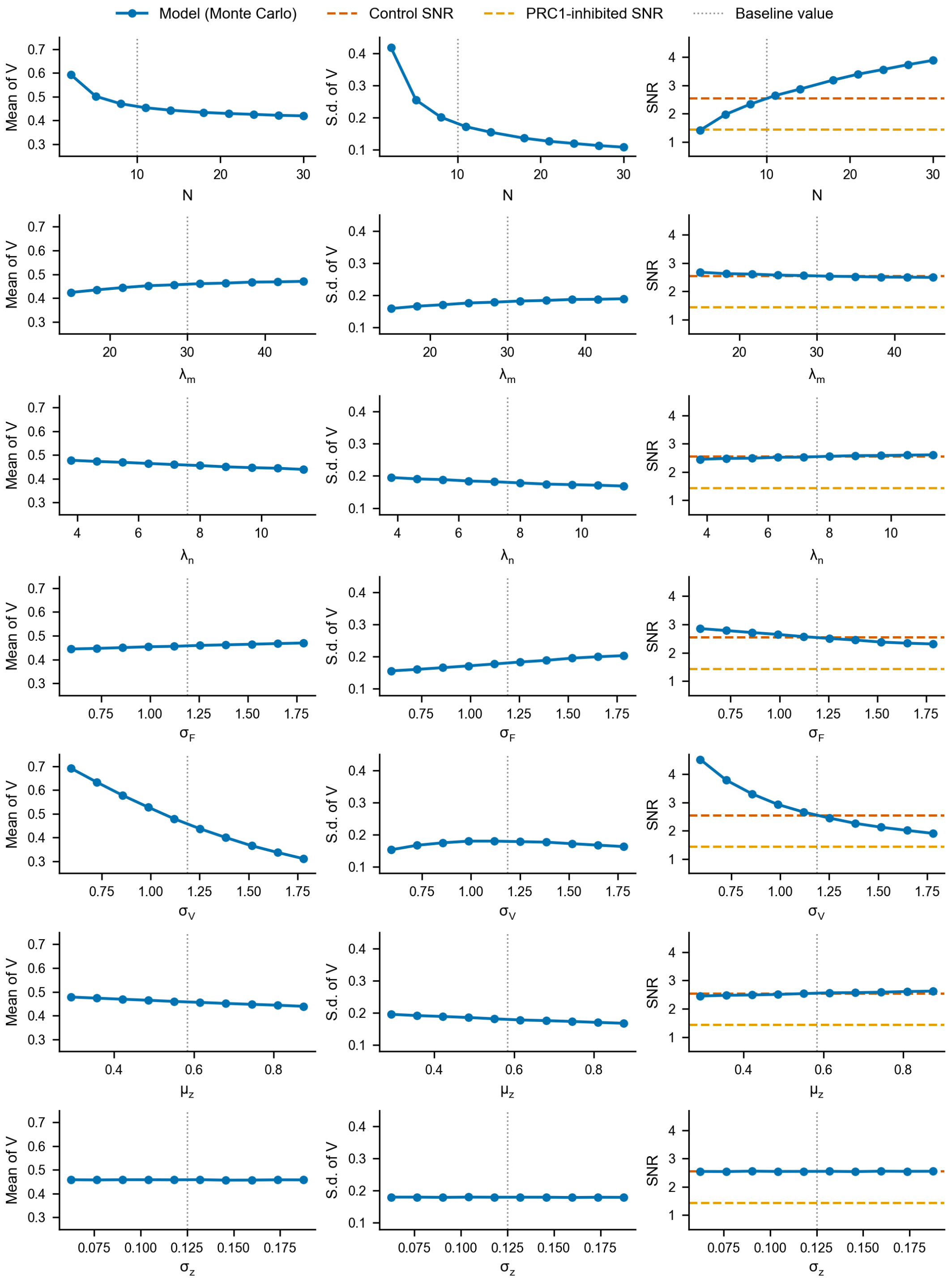


**Figure S3. Sensitivity analysis of the stochastic interpolar model.** In each row, one parameter is varied while all others are held fixed at their baseline values (Table S1; dotted vertical lines). From top to bottom, the varied parameter is $N$ (integer values from 2 to 30), then $\lambda_m$, $\lambda_n$, $\sigma_F$, $\sigma_V$, $\mu_z$ and $\sigma_z$ (10 evenly spaced values from 0.5 to 1.5 times the baseline). The columns show the mean (left) and standard deviation (middle) of the sliding velocity $V$ (Eq. (S3)), in units of $V_0$, and its SNR (right). Each point is computed from $10^5$ Monte Carlo realizations. Dashed lines in the right column mark the measured SNR in control (vermilion, 2.55) and PRC1-inhibited (orange, 1.44) spindles (main-text Fig. 5B).
